# Risk Assessment and Monitoring of Certain Pollutants in Natural Honeys Imported from Different Countries and Marketing in KSA

**DOI:** 10.1101/2023.03.27.534451

**Authors:** Khaled A. Osman, Hala H. Elsayed Mohamed, Maher S. Salama

## Abstract

The physicochemical and antimicrobial properties, as well as metals and pesticide residue contents in honeys imported from different countries and marketing in KSA were investigated. The results indicated that the chemical composition of some of these honeys violated the most national and international guidelines. Also, honey samples showed greater antimicrobial activities against all the osmophilic microorganisms especially *Aspergillus flavus*. Pb and Cd were not detected in all the tested samples, while Cu levels were below the guideline value of 5 mg/kg. However, Zn, Fe, and Ni levels in most the tested samples did not comply with the legislation values of 5, 15, and 0.01-1.00 mg/kg, respectively, and may pose a health risk to consumers. Also, Mn was found in higher levels which can be attributed either to the production stages of honey or the region from where the honey has been taken. Regarding the pesticide residues, some residues were detected in honeys imported from Germany, Australia, and Turkey; however, the hazardous indices for all the detected residues were less than one, indicating that these residues could not pose a health risk. It can be concluded that natural honeys should be free of any objectionable metals and pesticides.

## 1. Introduction

Natural honeys are a natural sweet and complex substance, consist of combined nectar blossoms or the secretion of the living part of plants with specific substances of excreted by honeys (Codex Alimentarius Commission 2001; EU 2002) that can be used by humans without any processing (Iglesias et al. 2004) because they have an excellent nutritional value (Song et al. 2018; Hu et al. 2021; Wang et al. 2021). Honeys contain sugars, proteins, and trace elements that are playing an important role in a number of biochemical processes in humans (Hak-Gil et al. 1988; Falco et al. 2003; Garcia et al. 2005). The chemical composition of these honeys depends on the type of plants and the climate conditions in which the plants grow (Abou-Tarboush et al. 1993; Anklam 1998).

Pesticides are widely used either in agriculture and public health sector to control a variety of pests and vector-born-diseases. During foraging periods especially near agricultural areas, bees exposed to a variety of pesticides causing bees fed on blossom contaminated with pesticide residues (Lazarus et al. 2021). Many studies were investigated to monitor residues of pesticides in natural honey collected from different countries (Menkissoglu-Spiroundi et al. 2001; Malhat et al. 2019; Osman et al. 2021; Mahdavi et al. 2022). Although the maximum residue limits of some pesticides (MRLs) in honey been established (EC 1999; FDA 2003), others are not included in the Codex Alimentarius (1998). Because of the possible risks from honey consumption, setting regulation threshold limits for the either heavy metals or pesticides should be considered (FDA 2003; Scivicco et al. 2022).

Because Saudi Arabia (KSA) produces approximately 9,000 metric tons of honey annually which is too small to satisfy the local demand (Al-Ghamdi 2007), therefore, natural honeys are imported from different countries to face the increasing in population and consumption (Alnafissa and Alderiny 2020), where the imported natural honeys increased from 6400 to 16500 tons during the last two decades (Ministry of Environment, Water, and Agriculture 2018). Some of metals and pesticide residues are being toxic if they violate the safety levels (Codex Alimentaris Commission 1993; Osman et al. 2007; Osman et al. 2021), therefore, honey must be free of any objectionable contents because they are used by humans without any processing.

Because negligible data is available on the composition of honeys imported from the major suppliers of honey, namely, Australia, France, Germany, India and Turkey, therefore the present study aimed to determine the physicochemical properties as well as metals and residues of pesticide contents in these honeys. Furthermore, a risk assessment for contaminated honey consumption by humans was performed by compare the estimate daily intakes (EDIs) with the acceptable daily intakes (ADIs).

## 2. Material and Methods

Chemicals used in the present study are of analytical-reagent grade.

### 2.1. Reagents and solutions

Standards elements (1000 mg/l, J.B. Baker Inc., Phillipsburg, NJ, USA) of Fe, Zn, Cu, Mn, Pb, Cd, and Ni were used to prepare the working standard solutions in deionized water obtained from a (PURELAB Option-R, ELGA, UK). Analytical grades of nitric acid (65%) and hydrochloric acid (36%), and the HPLC grades of either ethyl acetate, hexane, acetonitrile, and isooctane were purchased from BDH Company (UK), while hydrogen peroxide (30 %) was from WINLAB (UK). Standard isobenzan was purchased from Dr. Ehrenstorfer GmbH (Augsburg, Germany), while of carbaryl, pirimicarb, tolclofos-methyl, and monocrotophos were obtained from the Environmental Protection Agency (EPA, USA) with certified purities of 99% for all the investigated pesticides.

### 2.2. Honey samples

Seventy-five honey samples imported from Australia, France, Germany, India and Turkey (15 samples for each country, 0.5-1 Kg each) were purchased from supermarkets located in KSA, immediately transferred to the laboratory and kept at 4 °C until analysis.

### 2.3. Physicochemical properties of honeys

Sugar, moisture, ash, pH, total acidity, electric conductivity (EC) and hydroxymethyl furfural (HMF) in tested samples were determined according to the methods of AOAC (2002) as previously described before (Osman et al. 2021). Protein content (mg/g fresh weight) was determined according to the method of Lowry et al. (1951), while the method of Louveaux et al. (1978) was used to determine the number of pollen grains (NPG), honeydew element (HDE) and botanical element (BEN) using the light microscope (Olympus, Japan). Reflectance of honey samples was measured against white background at wavelength ranging from 300 to 800 nm (Color Flex, Hunter Lab, USA) to measure the color of honeys.

### 2.4. The antimicrobial activity of honeys

The *in vitro* potency of the tested honeys to inhibit of *Penicillium spp*., *Aspergillus niger*, *Aspergillus flavus*, *Alternaria alternate*, *Staphylococcus spp*., and *Saccharomyces spp* was carried using the paper disk technique.

### 2.5. Determination of minerals in honey samples

The levels of metals (Fe, Zn, Cu, Mn, Pb, Cd, and Ni) in the tested honeys were determined according to the method of Hernández et al. (2005) using the atomic absorption spectrometry with operating conditions as previously conducted (Osman et al. 2021) with correction of the matrix interference. Working solutions ranged from 0.0-2.0 mg l^-1^ in case of all the tested elements except in case of Fe it ranged from 0.0-10.0 mg l^-1^ were prepared from metal standard solution (1000 mg/l). Recoveries were carried out at levels of 0, 0.25, and 0.5 mg g^-1^ and ranged from 90-102%. The recorded detection limits were found to be 8.00, 1.40, 0.37, 0.81, 0.60, 1.00, and 0.50 g/kg for Fe, Zn, Cu, Mn, Pb, Cd, and Ni, respectively.

### 2.6. Pesticide residue analysis

Because honey is a complex foodstuff contains high levels of beeswax, proteins, metals and several classes of pesticides that may affect the results of analysis, therefore the analysis of pesticide gas chromatography coupled with mass spectrometry (GC-MS) is highly sensitive and selective technique (Karazafiris et al. 2008; Tette et al. 2016). In the present study, a multiresidue method for pesticide analysis in honey were conducted using solid phase extraction technique for extraction and clean-up, analyzed by GC-MS. Retention times (Rt) and the presence of three fragment ions (m/z) were considered to identify the residues (Berrada et al. 2010).

The percentages of recovery precision of the method were carried out by adding 0.05, 0.10 and 0.50 mg/kg form each pesticide to honey samples free from pesticides extracted as previously described. Recoveries and precision ranged from 92-105% and 3-12%, respectively, which are acceptable (Osman et al. 2010, 2011 and 2021). The limits of detection (LOD) and quantitation (LOQ) were calculated and found to be 18, 10, 20, 12 and 5 μg/kg for LOD, while LOQ values were 60, 33, 67, 40 and 17 μg/kg for carbaryl, tolclofos-methyl, isobenzan, monocrotophos and pirimicarb, respectively.

### 2.7. Statistical analysis

All data were calculated as mean ± standard deviation (SD) and analyzed using analysis of variance technique (ANOVA) at significant level of 0.05.

## 3. Results and Discussion

### 3.1. Physicochemical properties of honeys

Water, sugar, ash, and protein contents, pH, total acidity and EC are usually used to characterize the properties of honeys (Campos et al. 2001; Osman et al. 2007 and 2021). Data in Table (1) the percentages of moisture contents ranged from 22.93-23.73%, with non-significant differences between all the samples imported from different countries, indicating bad degree of maturity for all samples and violated the specifications of <20% moisture (Anonymous 1992; EU 2002). Unfortunately, the highest percentage of moisture renders honey to ferment and spoil rapidly and affects the shelf-life of the honey (Pérez-Arquillué et al. 1994). This moisture depends on many factors such as weather, original moisture of the nectar, harvest season, the conditions of storage and the degree of honey maturity (Terrab et al. 2003b).

**Table 1.**
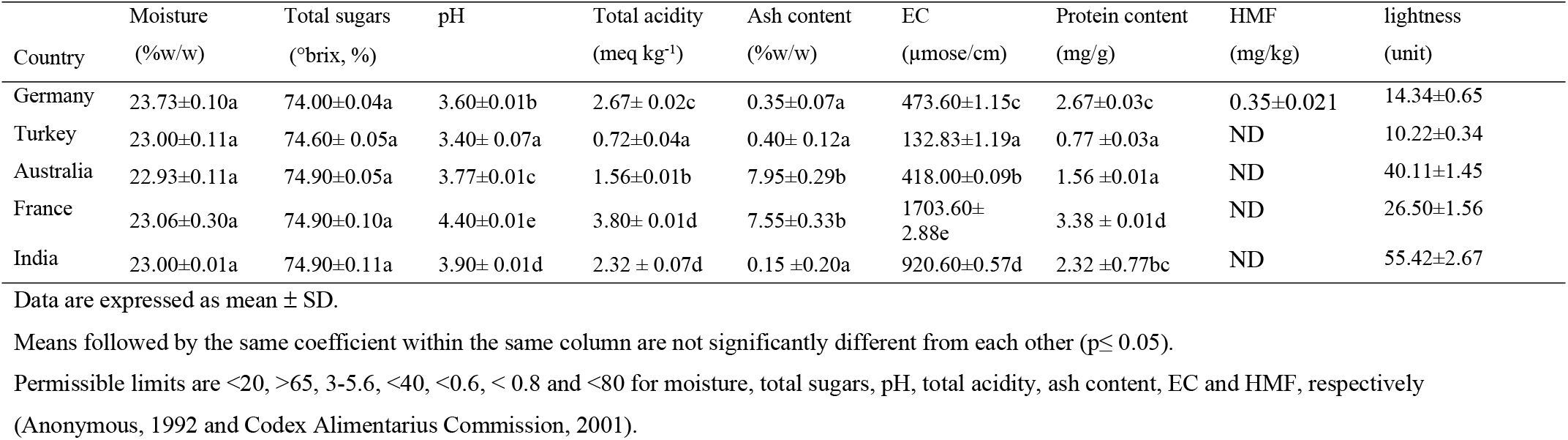
Physicochemical properties of imported honeys from different countries.

Also, data in Table (1) show that all the honey samples had high sugar contents of >72%, which are in agreement with the guidelines of > 65% of total sugars (Anonymous 1992; Codex Alimentarius Commission 2001; EU 2002). The obtained percentages of sugar are close to that found in different countries (Conti 2000; Sanz et al. 2005; Osman et al. 2021).

The pH values of all the tested samples ranged from 3.60-4.40 (Table 1) and within the limits established by the national and international specifications of 3.00-5.60 (Anonymous 1992; Codex Alimentarius Commission 2001; EU 2002). The obtained pH values are similar to that for floral honeys (White 1978) and suitable for the yeasts and molds growth (Conti et al. 1998).

Regarding the total acidity, none of the tested samples exceeded the limit of 40 meq/kg established (Anonymous 1992; Codex Alimentarius Commission 2001; EU 2002), where the values for the samples ranged from 0.72-3.80 meq/kg (Table 2). The variation in the total acidity among honeys imported from different countries may be attributed to either variation in the contents of organic and inorganic acids, the harvest season and/or floral types (El-Sherbiny and Risk 1979; Pérez-Arquillué et al. 1994). The pH of the honey is not directly related to free acidity because of the buffering action of the various acids and minerals present in honey (Abou-Tarboush et al. 1993).

**Table 2.** Quantitive pollen analysis of imported honeys from different countries

| Country | NPG<sup>1</sup> | HDE<sup>2</sup> | BEN<sup>3</sup> |
| --- | --- | --- | --- |
| Germany | 3.12×10<sup>4</sup> a | 5.72×10<sup>6</sup> c | 5.75×10<sup>6</sup> b |
| Turkey | 1.40×10<sup>6</sup> c | 8.04×10<sup>6</sup> d | 8.18×10<sup>6</sup> c |
| Australia | 0 | 5.70×10<sup>6</sup> b | 5.72×10<sup>6</sup> b |
| France | 0 | 2.41×10<sup>8</sup> a | 2.41×10<sup>8</sup> d |
| India | 1.56×10<sup>5</sup> b | 3.40×10<sup>5</sup> a | 4.96×10<sup>5</sup> a |

<sup>1</sup> number of pollen grains; <sup>2</sup> number of honeydew indicator elements; <sup>3</sup> number of botanical elements (NPG + HDE).
Means followed by the same coefficient within the same column are not significantly different from each other ( $p \leq 0.05$ ).

**Table 3.** Antimicrobial effects of honeys imported from different countries.

| Source | % of inhibition |  |  |  |  |  |
| --- | --- | --- | --- | --- | --- | --- |
|  | *Pencillium spp.* | *A. alternate* | *A. niger* | *A. flavus* | *Staphylococcus spp.* | *Saccaromyce spp* |
| Germany | 42.31±1.00a | 50.00 ±1.0c | 42.30 ±0.70b | 100.00±1.40a | 46.16 ±1.50a | 42.39 ±1.10a |
| Turkey | 100.00 ±1.00a | 100.00±1.0d | 100.00±2.00f | 100.00±1.60a | 42.31 ±1.52a | 42.31 ± 1.00a |
| Australia | 100.00±1.05a | 42.00 ±1.0a | 100.00±2.00f | 100.00±1.00a | 46.20 ±1.10a | 42.11 ± 1.05a |
| France | 100.00±1.01a | 46.20 ±1.0b | 42.31 ±1.00a | 100.00±1.00a | 46.66 ±1.30a | 42.81 ±1. 04a |
| India | 42.50±0.90a | 42.55 ±1.0a | 42.65 ±1.00a | 100.00±1.00a | 46.00 ±1.40a | 42.66 ± 1.00a |

Mean is expressed as % of control (n= 3).
Data are expressed as mean ± SD. Means followed by the same coefficient within the same column are not significantly different from each other ( $p \leq 0.05$ ).

The ash contents in honey samples imported from either Australia or France found to be 7.95 and 7.55%, respectively, and violated the limit allowed of 0.6% (Anonymous 1992; Codex Alimentarius Commission 2001; EU 2002) (Table 1). Sometimes, beekeepers are forced to feed bees on sucrose, for maintaining healthy bee colonies, and this may explain the increasing the ash contents in honeys imported from some country. However, the ash contents of samples imported from Germany, Turkey and India were within the allowed limit. Natural honeys normally have low ash contents which influenced by the botanical origin and the technique used for the determination (Abou-Tarboush et al. 1993).

The highest EC recorded in honeys imported either from France (1703.60 μmoose/cm) or India (920.60 μmoose/cm) confirmed the high content of mineral substances in flower honey, and did not meet the requirement of 800 μmoose/cm (Table 2). Also, the EC test may provide evidence for honey adulteration due to honeybees overfeeding with sucrose-syrup solutions (Nisbet et al. 2018). However, honey imported from Germany, Turkey and Australia was found to be within the allowed limit. The content of elements in honeys significantly correlates with the EC, where the higher contents of minerals, the higher values of EC (Bogdanov et al. 2007).

Also, the data show that all the honey samples are rich in protein content because contents were >1.0 mg/g and a had a pollen predominance of the same floral origin (Azeredo et al. 2003), except samples imported from Turkey contained 0.77 mg/g honey. The major components of pollen are proteins, amino acid, lipids and sugars (Atrouse et al. 2004).

Data in Table (2) illustrate that HMF was not detected in all the investigated honeys except in samples collected from Germany contained 0.355 ppm which is lower than the maximum allowed of 80 ppm without any coloration (Anonymous 1992; Codex Alimentarius Commission 2001). This means that all samples were not adulterated with commercial sugar or exposed to high temperatures and harvested after a short time (Azeredo et al. 2003; Khan et al. 2005).

The color of tested samples ranged from amber (14.34-26.50 units) to dark amber (40.11-55.42 units), where the dark samples have antioxidant properties (Stephan, 1989). The lightness values found in the present study are close to that found in Moroccan (Terrab et al. 2003a) and Saudi honeys (Osman et al. 2007; Osman et al. 2021). The differences in color of honey are due to the botanical origin and the number of suspended particulates such as pollen (Atrouse et al. 2004).

### 3.2. Quantitive pollen analysis of imported honey samples

The results of the microscopic analysis of honeys illustrated that the number of NPG, HDE and BEN in 10 g of honey sample ranged from 0 (Australia and France) - 1.56×10^5^ (India), 3.40×10^5^ (India) - 8.04×10^6^ (Turkey) and 4.96×10^5^ (India) - 8.18×10^6^ (Turkey), respectively (Table 2). These honeys are considered as rich in their NPG contents (Maurizio 1979) except that imported from Australia and France, while honeys collected from France had the highest counts either of HDE or BEN, followed by Turkey, Germany, Australia, and then India. The number of pollen grains recorded in samples collected from different countries may be due to the differences in the origin of plants that the bees fed.

### 3.3. Antimicrobial activities of honeys

The antimicrobial activities of honeys against the tested osmophilic microorganisms illustrated that all the tested samples had high potency to inhibit these microorganisms with inhibition percentages of 42-100% and *A. flavus* was the most sensitive one followed by *Pencillium spp*., *A. niger*, *A. alternate*, *Staphylococcus spp*., and then *Saccaromyce spp*. Also, the results revealed that honeys imported from Turkey were found to be the most potent against these osmophilic microorganisms followed by that imported from Australia and then France, and this antimicrobial effect of honeys may due to their contents of flavonoids, phenols, and organic acid, as well as high osmotic pressure, high acidity, and production of hydrogen peroxide (Li et al. 2001; Malika et al. 2004; Iurlina and Fritz 2005). These properties render natural honeys are unsuitable to support the growth of microorganisms and rarely to spoil during storage at home.

### 3.4. Metals analysis

Data in Table (4) illustrate that samples imported from India was found to contain the highest levels of Fe, Cu, and Mn with levels of 10.88, 3.55, and 6.79 mg kg^-1^, respectively, while samples imported from Australia and Germany contained the highest levels of Zn (5.92 mg kg^-1^) and Ni (1.75 mg kg^-1^), respectively. The levels of Fe, Zn and Cu in all samples were lower than the guideline values of 15, 5, and 5 mg/kg (Codex Alimentarius Commission 2001), respectively, except that samples collected from Australia contained levels of Zn higher than the established guideline. The obtained Zn and Fe levels were lower than levels found honeys from KSA (Osman et al. 2021) and Turkey (Tuzen et al. 2007).

**Table 4.**
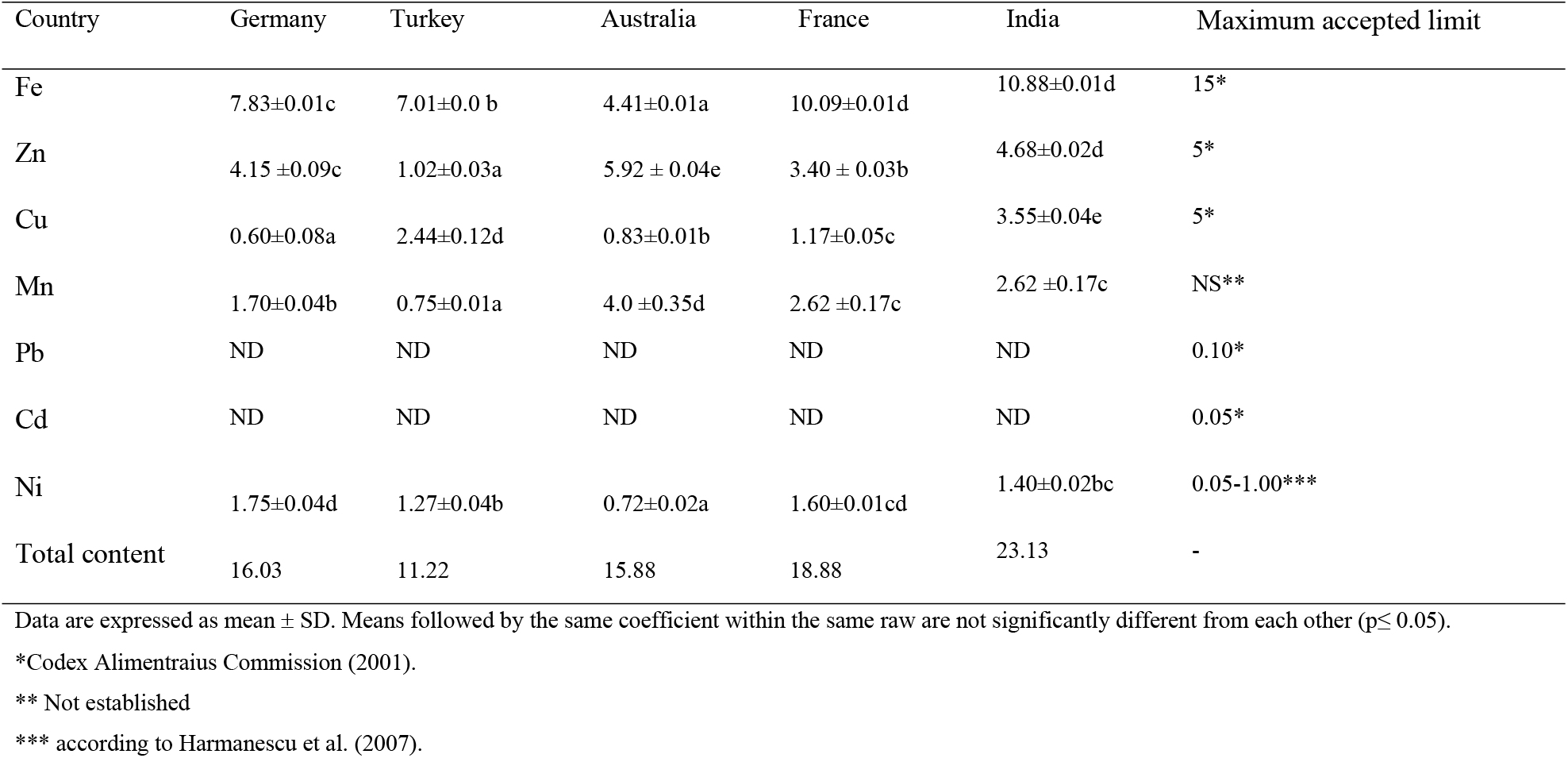
Levels (mg kg^-1^ fresh weight) of elements in natural honeys imported from different countries.

| Country | Germany | Turkey | Australia | France | India | Maximum accepted limit |
| --- | --- | --- | --- | --- | --- | --- |
| Fe | 7.83±0.01c | 7.01±0.0 b | 4.41±0.01a | 10.09±0.01d | 10.88±0.01d | 15* |
| Zn | 4.15 ±0.09c | 1.02±0.03a | 5.92 ± 0.04e | 3.40 ± 0.03b | 4.68±0.02d | 5* |
| Cu | 0.60±0.08a | 2.44±0.12d | 0.83±0.01b | 1.17±0.05c | 3.55±0.04e | 5* |
| Mn | 1.70±0.04b | 0.75±0.01a | 4.0 ±0.35d | 2.62 ±0.17c | 2.62 ±0.17c | NS** |
| Pb | ND | ND | ND | ND | ND | 0.10* |
| Cd | ND | ND | ND | ND | ND | 0.05* |
| Ni | 1.75±0.04d | 1.27±0.04b | 0.72±0.02a | 1.60±0.01cd | 1.40±0.02bc | 0.05-1.00*** |
| Total content | 16.03 | 11.22 | 15.88 | 18.88 | 23.13 | - |

Data are expressed as mean ± SD. Means followed by the same coefficient within the same raw are not significantly different from each other ( $p \leq 0.05$ ).
\*Codex Alimentarius Commission (2001).
\*\* Not established
\*\*\* according to Harmanescu et al. (2007).

Also, the data illustrated that all imported honey samples were found to be free of either Pb or Cd (Table 4). Honey samples collected from many apiaries located in KSA were also found free of Cd, while had a significant Pb contents (Osman et al. 2021). Also, honeys collected Malaysia (Shahid et al. 1987), Hungary (Ajtony et al. 2007), Algeria (Di Bella et al. 2022) were found to have a significant Pb levels. The levels of Mn found in the present study were higher than that found in honey samples collected from KSA (Osman et al. 2021), and similar to that reported by many investigators (Conti 2000; Devillers et al. 2002; Ioannidou et al. 2005). Unfortunately, the guideline value has not been published for Mn. The higher levels of Mn can be attributed either to the production stages of honey and/or the region from which the honey has been taken (Ioannidou et al. 2005). Regarding the levels of Ni, it was found that samples imported from the all the countries did not comply the legislation levels of 0.01-1.00 mgkg^-1^ (Harmanescu et al. 2007) except those honeys imported from Australia (Table 4). These levels were less than that in honeys either from Turkey (Tuzen et al. 2007) or KSA (Osman et al. 2021). In the present study, the mineral contents ranged from 11.22 in honeys imported from Turkey to 23.13 mg kg^-1^ in honeys imported from India which within the accepted range (Hernández et al. 2005; Solayman et al. 2016).

### 3.5. Pesticide residues

Carbaryl, pirimicarb, and tolclofos-methyl at levels of 0.55, 0.017, and 0.054 mg kg^-1^ were detected in samples imported from Germany, while samples imported from Australia contained carbaryl and isobenzan at levels of 17.08 and 82.75 mg kg^-1^, respectively (Table 5). Also, honeys imported from Turkey were found to contain monocrotophos at level of 2.78 mg kg^-1^. All the detected pesticides are recommended to be used to protects crops and can be introduced into hives by workers bees (Anju et al. 1997; Mullin et al. 2010; Chauzat et al. 2011, Kasiotis 2014). Different pesticides from different chemical groups were detected in honeys collected from different countries (Calatayud-Vernich et al. 2016; Fulton 2019; Gawel et al. 2019; Lazarus et al. 2021; Osman et al. 2021). These residues in foodstuffs are generally legislated to reduce the exposure of humans to unnecessary intakes of these pesticides. Unfortunately, none of the detected pesticides in the present study included in the Codex Alimentarius (1998). However, the maximum residues limits (MRLs) for amitraz, coumaphos, and cyamizole, and fluvalinate have been regulated (EC 1990; FDA 2003). From a potential health perspective, it is important to compare the estimate daily intakes (EDIs) calculated according to the guidelines (WHO 1997; FAO 2002) with the established the amount of that pesticide that can be ingested daily by a human being during an entire lifetime without an appreciable risk to the health (the acceptable daily intakes, ADIs). The EDI was calculated based on the following equation:

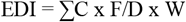

where C, F, D, and W represent the sum of concentration of pesticide in each commodity (mg/kg), the mean annual intake of imported honey per person, the number of days in a year (365), and the mean body weight of human (60 kg), respectively.

**Table 5.** Levels (mg kg^-1^ fresh weight) of pesticide residues detected in natural honeys imported from different countries and their estimated (EDIs) and accepted daily intakes (ADIs).

| Pesticide | Country |  |  |  |  | ADI<br>(mg/kg body<br>weight/day)<sup>2</sup> | EDI<br>(mg/kg body<br>weight/day) | Hazard index<br>(EDI/ADI, %) |
| --- | --- | --- | --- | --- | --- | --- | --- | --- |
|  | Germany | Turkey | Australia | France | India |  |  |  |
| Carbaryl | 0.55±0.05a | ND | 17.08±0.23b | ND | ND | 7.50 | 4.07x10<sup>-4</sup> | 5.43x10<sup>-3</sup> |
| Pirimicarb | 0.017±0.01 | ND | ND | ND | ND | 35.0 | 3.93x10<sup>-7</sup> | 1.12x10<sup>-6</sup> |
| Tolclofos-methyl | 0.054±0.01 | ND | ND | ND | ND | 64.0 | 1.25x10<sup>-6</sup> | 1.95x10<sup>-5</sup> |
| Monocrotophos | ND<sup>1</sup> | 2.78±0.14 | ND | ND | ND | 0.60 | 6.42x10<sup>-5</sup> | 1.07x10<sup>-2</sup> |
| Isobenzan | ND | ND | 82.75±0.82 | ND | ND | NS<sup>3</sup> | 1.91x10<sup>-3</sup> | - |

Data are expressed as mean ± SD. Means followed by the same coefficient within each column do not significantly different from each other (p≤ 0.05).
<sup>1</sup>ND means not detected
<sup>2</sup>According to Codex Alimentarius Commission (FAO/WHO, 2004).
<sup>3</sup> NS means ADI has not been established.

In 2017, KSA imported about 16500 tons of honeys from different countries with a population of 32.60 million at the same year (Ministry of Environment, Water, & Agriculture, 2018) and average annual consumption of imported honey/person being 0.506 kg. Data in Table (5) compare the EDIs contribution of honey consumed to the intake of these pesticides with ADIs (FAO/WHO, 2004). Data in Table (5) illustrate that isobenzan had the highest value of EDI followed by carbaryl, monocrotophos, tolclofos-methyl, and then pirimicarb, where the EDIs ranged between 3.93×10^-7^ to 1.91×10^-3^ mg/kg body weight/day. The obtained EDIs of the tested pesticides are much lower than the ADIs, illustrating that the consuming of imported natural honeys has a minimal contribution to the toxicological risk. No ADI for isobenzan has been established. The obtained hazard indices, HI (EDI/ADI) were 1.07×10^-2^, 5.43×10^-3^, 1.95×10^-5^, and 1.12×10^-6^ for monocrotophos, carbaryl, tolclofos-methyl, and pirimicarb, respectively. The consumption of these honeys could not pose health risk for KSA population because HI values < 1 (Darko and Akoto 2008) and similar to honeys collected from Iran (Mahdavi et al. 2022). However, honeys imported from Australia and India in the present should be considered due to their high levels of residues.

## 6. Conclusion

The chemical composition of some honey samples imported by KSA from different countries do not comply all the national and international guideline values and contained high levels of metals such as Zn and Ni and different pesticide residues belong to different chemical groups such as carbaryl and pirimicarb (carbamates), tolclofos-methyl and monocrotophos (organophosphates) and isobenzan (organochlorines) which may pose a health problem especially for human who consume honey as a source of useful food on a regular basis (Altunay et al. 2019; Osman et al. 2021). Because honey is a universal food with the high demands on the quality of imported honeys among consumers, therefore the assurance of its components safety via monitoring programs is critical.

## Compliance with ethical standards

### Data Availability

The data presented in this study are available on request from the corresponding author. The data are not publicly available due to privacy restrictions.

### Consent to participate

The authors declare that they have no known competing financial interests or personal relationships that could have appeared to influence the work reported in this paper.

### Consent to publish

Informed consent was obtained from all individual participants included in the study.

### Plant Reproducibility

This article does not contain any studies with Plant Reproducibility performed by any of the authors.

### Clinical trials registration

This study does not contain any studies with clinical trials performed by any of the authors.

### Funding

There are currently no Funding Sources in the list.

## References

Abou-Tarboush H, Al-Kahtani H, El-Sarrage M (1993) Floral type identification and quality evaluation of some honey types. Food Chem 46:13–17.

Ajtony Z, Bencs L, Haraszi R, Szigeti J, Szoboszlai N (2007) Study on the simultaneous determination of some essential and toxic trace elements in honey by multi-element graphite furnace atomic absorption spectrometry. Talanta 71:683–90.

Al-Ghamdi AA (2007). Beekeeping and Honey Production in Saudi Arabia. 5th Conference of the Arab Beekeepers Association, Tripoli.

Al-Khalifa AS, Al-Arify IA (1999) Physicochemical characteristics and pollen spectrum of some Saudi honey. Food Chem 67:21–25.

Alnafissa M, Alderiny M (2020) Analysis of Saudi demand for imported honey using an Almost Ideal Demand System (AIDS). J Saudi Soci Agri Sci 19:293–298.

Altunay N, Elik A, Gürkan R (2019) Monitoring of some trace metals in honeys by flame atomic absorption spectrometry after ultrasound assisted-dispersive liquid liquid microextraction using natural deep eutectic solvent. Microchem J 147:49–59.

Anju R, Beena K, Gahlawat SK, Sihag RC, Kathpal TS (1997) Multiresidue analysis of market honey samples for pesticidal contamination. Pestic Res J 9:226–230.

Anklam E (1998) A review of the analytical methods to determine the geographical and botanical origin of honey. Food Chem 63(4):549–562.

Anonymous (1992) Honeybees in Council of Gulf Country. Journal of Standards, Riyadh, Part 1, Vo. 21(In Arabic).

AOAC (2002) Official Methods of Analysis. 17th (Ed), Association of Official Analytical Chemists Gaithersburg, Maryland, USA.

Atrouse OM, Oran SA, Al-Abbadi SY (2004) Chemical analysis and identification of pollen grains from different Jordanian honey samples. Inter J Food Sci Techn 39:413–417.

Azeredo L da C, Azeredo MAA, de Souza SR, Dutra VML (2003) Protein contents and physicochemical properties in honey samples of Apis mellifera of different floral origins. Food Chem 80:249–254.

Berrada H, Fernández M, Ruiz MJ, Moltó JC, Mañes J, Font G (2010) Surveillance of pesticide residues in fruits from Valencia during twenty months (2004/05). Food Cont 21:36–44.

Bogdanov S, Haldimann M, Luginbühl W, Gallmann P (2007) Minerals in honey: environmental, geographical and botanical aspects. J Apicul Res Bee World 46(4):269–275.

Calatayud-Vernich P, Calatayud F, Simó E, Suarez-Varela MM, Picó Y (2016). Influence of pesticide use in fruit orchards during blooming on honeybee mortality in 4 experimental apiaries. Sci Total Environm 54133–54141.

Campos G, della Modesta RC, da Silva TJP, Raslan DS (2001) Comparison of some components between floral honey and honeydew honey. Rivista do Instituto Adolfo Lutz 60:59–64.

Celli G, Maccagnani B (2003) Honey bees as bioindicators of environmental pollution. Bull Insect 56(1):137–139.

Chauzat MP, Martel AC, Cougoule N, Porta P, Lachaize J, Zeggane S, Aubert M, Carpentier P, Faucon JP (2011) An assessment of honeybee colony matrices, Apis mellifera (Hymenoptera: Apidae) to monitor pesticide presence in continental France. Environm Toxicol Chem 30:103–111.

Codex Alimentaris Commission (1993) Standard for Honey, Ref. no. CL 1993/14, SH, Codex Alimentarius Commission, FAO/WHO, Rome.

Codex Alimentarius (1998) Draft revised for honey at step 6 of the Codex Procedure. CX 5/10.2, CL 1998/12-S.

Codex Alimentarius Commission (2001) Revised Codex Standard for Honey, Codex STAN 12-1981, Rev.1 (1987), Rev.2.(2001)

Codex Alimentarius Commission (2001) Draft revised standard for honey. Alinorm 25:19–26.

Conti ME (2000) Lazio region (central Italy) honey: a survey of mineral content and typical quality parameters. Food Chem 11:459–463.

Conti ME, Saccares S, Cubadda F, Cavallina R, Tenoglio CA, Ciprotti L (1998) II Miele nel Lazio: Indagine sul contenuto in metallic in trace e radionuclide. Rivisita di Scienza dell Alimentazione 2:107–109.

Costat (1986) Version 2, Cohort Software. Minneapolis, MN, USA.

Crane E (1977) Constituents and characterization of Honey. In: A Book of Honey. Oxford University Press, London. p. 39.

Darko G, Akoto O (2008) Dietary intake of organophosphorus pesticide residues through vegetables from Kumasi, Ghana. Food Chem Toxicol 46:3703–3706.

Devillers J, Doré JC, Marenco M, Poirier-Duchen F, Galand N, Viel C (2002) Chemometrical analysis of 18 metallic and nonmetallic elements found in honeys sold in France. J Agri Food Chem 50(21):5998–6007.

Di Bella G, Licata P, Potortì AG, Crupi R, Nava V, Qada B, Rando R, Bartolomeo G, Dugo G, Lo Turco V. Mineral content and physicochemical parameters of honey from North regions of Algeria. Natural Product Research: 36(2):636–643.

EC (1999) Commission Regulation No. 2377/90 of 26 June 1990 laying down a Community Procedure for the stablemen of maximum residue limits of veterinary medicinal products in foodstuff of animal origin (as amended by regulations) ECC No. 2034/96 (OJL272 25.10.1996, p 2), No2686/98 (OJ L337 12.12.1998, p 20) No. 1931/99 (OJ L240 10.09.1999, p 3), and No. 239/99(OJ L 290 12.11.1999, p 5).

El-Sherbiny GA, Risk SS (1979) Chemical of both clover and cotton honey produced in A.R.E. Egypt J Food Sci 7:69–75.

EU (The Council of European Union) (2002) Council Directive 2001/110/EC of 20 December 2001 relating to honey. Offic J Eur Comm L10:47–52.

Falco G, Gomez-Catalan C, Llobet JM, Domingo JL (2003) Contribution of medicinal plants to the dietary intake of various toxic elements in Catalonia, Spain. Trace Elem Elect 20:120–124.

Falqui-Cao C, Wang Z, Urruty L, Pommier J-J, Montury M (2001) Focused microwave assistance for extracting pesticide residues from strawberries into water before their determination by SPME/ HPLC/DAD. J Agr Food Chem 49:5092–5097.

FAO (Food and Agriculture Organization) (2002) Submission and evaluation of pesticide residues data for the estimation of maximum residue levels in food and feed (1st ed). Rome: Food and Agriculture Organization. <http://www.fao.org/ag/AGP/AGPP/Pesticid/p.htm>.

FAO/WHO (Food and Agriculture Organization/World Health Organization). (2004). Food standards programme. Codex Alimentarius Commission. Twenty-seventh Session, Geneva, Switzerland.

FDA (Food and Drug Administration of the United States) (2003) Pesticide tolerances http://www.cfsan.fda.gov.

Fernández-Muiño MA, Sancho MT, Simal-Gándara J, Creus Vidal JM, Huidobro JF, Simal-Lozano J (1997) Acaricide residues in honey from Galicia (N. W. Spain). J Food Protect 60:78–80.

Fulton CA, Huff Hartz KE, Fell RD, Brewster CC, Reeve JD, Lydy MJ (2019). An assessment of pesticide exposures and land use of honey bees in Virginia. Chemosphere 222:489–493.

Garcia JCR, Garcia JB, Latorre CH, Martin SG, Crecent RMP (2005) Comparison of palladium-magnesium nitrate and ammonium dihydrogen phosphate modifiers for lead determination in honey by electrothermal atomic absorption spectrometry. Food Chem 91:435–439.

Gawel M, Kiljanek T, Niewiadowska A, Semeniuk S, Goliszek M, Burek O, Posyniak A (2019) Determination of neonicotinoids and 199 other pesticide residues in honey by liquid and gas chromatography coupled with tandem mass spectrometry. Food Chem 282(1):36–47.

Hak-Gil C, Myung-Kyoo H, Jae-Gil K (1988) The Chemical composition of Korean Honey. Korean J Food Sci Techn 20, 631–636.

Hamza AMT, Hassan AAK, El-Sarrage MS (1993) Floral-type identification and quality evaluation of some honey types. J Food Chem 46:13–17.

Harmanescu M, Popovici D, Gergen I (2007) Mineral micronutrients composition of bee’s pollen. Journal of Agroalimentary Processes Technol 13(1): 175–182.

Heather P (2012) Pesticides and Honey Bees: state of the sciences. Pan North America.

Human H, Archer C R, Du R, E. E, Pirk C W W, Nicolson SW (2014) Resistance of developing honeybee larvae during chronic exposure to dietary nicotine. J Insect Physiol 69:74–79.

Hernández OM, Fraga JMG, Jiménez AI, Jiménez F, Arias JJ (2005) Chracterization of honey from Canary Islands: Determination of mineral content by atomic absorption spectrometry. Food Chem 93:449–458.

Hu X, Zhang P, Du W, Jiang J, Chen X, Liu Y, Zhang Z, Tang B, Li P (2021) AIEgens enabled ultrasensitive point-of-care test for multiple targets of food safety: aflatoxin B1 and cyclopiazonic acid as an example. Biosen Bioelect 182:113188.

Iglesias MT, De Lorenzo C, Del Carmen Polo M, Martín-Ávarez PJ, Pueyo E. (2004) Usefulness honeydew and floral honey. Application to honeys from a small geographic area. J Agri Food Chem 52:84–89.

Ioannidou MD, Zachariadis GA, Anthemidis AN, Stratis JA (2005) Direct determination of toxic trace metals in honey and sugars using inductively coupled plasma atomic emission spectrometry. Talanta 65:92–97.

Iurlina MO, Fritz R (2005) Characterization of microorganisms in Argentinean honeys from different sources. Inter J Food Microbiol 105(3):297–304.

Karazafiris E, Tananaki C, Menkissoglu-Spiroudi U, Thrasyvoulou A (2008) Residue distribution of the acaricide coumaphos in honey following application of a new slow-release formulation. Pest Manag Sci 64(2), 165–171.

Kasiotis KM, Anagnostopoulos C, Anastasiadou P, Machera K (2014) Pesticide residues in honeybees, honey and bee pollen by LCMS/MS screening: Reported death incidents in honeybees. Sci Total Environ 485-486:633–642.

Khan M, Qaiser NM, Raza SM, Rehman M (2005) Physicochemical properties and pollen spectrum of imported and local samples of blossom honey from the Indian market. Inter J Food Sci Tech 40:1–8.

Lazarus M, Lovakovic BT, Orct T, Sekovanic A, Bilandzic N, Dokic M, Kolanovic BS, Varenina I, Juric A, Lugomer DM, Bubalo B (2021) Difference in pesticides, trace metal(loid)s and drug residues between certified organic and conventional honeys from Croatia. Chemosphere 266:128954.

Li CQ, Pignatelli B, Ohshima H (2001) Increased oxidative and nitrative stress in human stomach associated with cag A +Helicobacter pylori infection and inflammation. Diges Dis Sci, 46:836–844.

Li Y, Kelley RA, Anderson TD, Lydy MJ (2015) Development and comparison of two multi residue methods for the analysis of select pesticides in honey bees, pollen, and wax by gas chromatography–quadrupole mass spectrometry. Talanta 140:81–87.

Louveaux J, Maurizio A, Vorwohl G (1978) Methods of melissopalybology, Bee World 59:139–157.

Lowry OH, Rosebrough NJ, Farr WL, Randall RJ (1951) Protein measurement with folin phenol reagent. J Biol Chem 193(1):265–275.

Mahdavi V, Eslami Z, Omidvari Z, Rezadoost H, Thai VN, Fakhri Y (2022) Carcinogenic and non-carcinogenic risk assessment induced by pesticide residues in honey of Iran based on Monte Carlo simulation. J Food Comp Anal 109:104521.

Malhat FM, Haggag MN, Loutfy NM, Osman MA., Ahmed MT (2015) Residues of organochlorine and synthetic pyrethroid pesticides in honey, an indicator of ambient environment, a pilot study. Chemosphere 120:457–461.

Malika N, Mohamed F, Adlouni C El-A (2004) Antimicrobial activities of natural honey from aromatic and medicinal plants on antibioresistant strains of bacteria. Inter J Agri Biol 6:289–293.

Maurizio A (1979) Microscopy of Honey. In: Crane E (Ed), Honey. A Comprehension Survey, pp. 240–257, London: Heinemann.

Menkissoglu-Spiroundi U, Tsigouri AD, Diamantidis GC, Thrasyvoulou AT (2001) Residues in honey and beeswax caused by beekeeping treatments. Fres Environ Bull 5:445–450.

Ministry of Agriculture of Saudi Arabia. (2006) Saudi Arabian food balance sheets for the period (2002-2004) to (1999-2001) (6 ed.). Ministry of Agriculture, Affair of Research and Agricultural Development-Organization of Studies, Planning and Statistics.

Ministry of Environment Water and Agriculture (2018) Open data Library, Information and Statistics. Saudi Arabia, Riyadh. <https://www.mewa.gov.sa/en.<

Mullin CA, Frazier M, Frazier JL, Ashcraft S, Simonds R, Van Engelsdorp D, Pettis JS (2010) High levels of miticides and agrochemicals in North American apiaries: Implications for honey bee health. PLoS One 5: e9754.

Nisbet C, Kazak F, Ardal Y (2018) Determination of quality criteria that allow differentiation between honey adulterated with sugar and pure honey. Biol Trace Element Res 186:288–293.

Osman KA, Al-Doghairi MA, Al-Otaib ND (2021) Spatial distribution of environmental pollutants in natural honeys collected from some regions of Saudi Arabia. Journal of Apicultural Research, 60(1), 188–197, https://doi.org/10.1080/00218839.2020.1727658.

Osman KA, Al-Doghairi MA, Al-Rehiayani SM, Helal MID (2007) Mineral content and physicochemical properties of natural honeys produced in Al-Qassim region, Saudi Arabia. J Food Agri Environ 5 (3&4):142–146.

Osman KA, Al-Humaid AM, Al-Rehiayani SM, Al-Redhaiman KN (2010). Monitoring of Pesticide Residues in Vegetables Marketed in Al-Qassim Region, Saudi Arabia. Ecotoxicol Environ Saf 73:1433–1439.

Osman KA, Al-Humaid AM, Al-Rehiayani SM, Al-Redhaiman KN (2011) Estimated daily intake of pesticide residues exposure by vegetables grown in greenhouses in Al-Qassim Region, Saudi Arabia. Food Cont 22:947–953.

Pérez-Arquillué C, Conchello P, Arino A, Juan T, Herresa A (1994) Quality evaluation of Spanish rosemary (*Rosomarinus officinalis*) honey. Food Chem 51:207–210.

Sanz ML, Gonzalez M, de Lorenzo C, Sanz J, Martínez-Castro I (2005). A contribution to the differentiation between nectar honey and honeydew honey. Food Chem 91:313–317.

Scivicco M, Squillante J, Velotto S, Esposito F, Cirillo T, Severino L (2022) Dietary exposure to heavy metals through polyfloral honey from Campania region (Italy). J Food Comp Anal 114:104748.

Serrano S, Villarejo M, Espejo R, Jordal M (2004). Chemical and physical parameters of Andalusian honey: classification of Citrus and Eucalyptus honeys by discriminant analysis. Food Chem 87:619–625.

Shahid SM, Sionge TE, Chong YH (1987) Lead content of some Malaysian foodstuffs. Asian Food 3:25–29.

Solayman M, Islam MA, Paul S, Ali Y, Khalil MI, Alam N, Gan SH (2016) Physicochemical properties, minerals, trace elements, and heavy metals in honey of different origins: A comprehensive review. Compreh Rev Food Sci Food Saf 15:219–233.

Song S, Zhang C, Chen Z, He F (2018) Simultaneous determination of neonicotinoid insecticides and insect growth regulators residues in honey using LC–MS/MS with anion exchanger disposable pipette extraction. J Chromatog A 1557:51–61.

Stephan B (1989) Determination of pinocembrin in honey using HPLC. J Api Res 28:55–57.

Sullivan A, Sheffrin SM (2003) Economics: Principles in Action. Pearson Prentice Hall, Upper Saddle River, 157.

Terrab A, Díez MJ, Heredia FJ (2003a) Palynological, physicochemical and colour characterization of Moroccan honeys. I. River red gum (*Eucalyptus camaldulensis* Dehnh) honey. Inter J Food Sci Techn 38, 379–386.

Terrab A, Díez MJ, Heredia FJ (2003b) Palynological, physicochemical and colour characterization of Moroccan honeys. II. Orange (Citrus sp.) honey. Inter J Food Sci Technol 38:387–394.

Tette PAS, Guidi LR, Glória MBA, Fernandes C (2016) Pesticides in honey: A review on chromatographic analytical methods. Talanta 149, 124–141.

Tuzen M, Silici S, Mendil D, Soylak M (2007) Trace element levels in honeys from different regions of Turkey. Food Chem 103:325–330.

USAID 2012 The World Market for Honey. http://www.ethiopia-ciafs.org/ciafs@fintrac.com/ www.fintrac.com/Market Survey #01/September 2012.

Wallner K (1999) Varroacides and their residues in bee products. Apidologie 30:235–248.

Wang Z, Ma S, Sun B, Wang F, Huang J, Wang X, Bao Q (2021) Effects of thermal properties and behavior of wheat starch and gluten on their interaction: A review. Inter J Biol Macromol 177, 474–484.

White Jr JW (1978) Honey. Adv Food Res 24:287–374.

White JW, Crane E (1979) A Comprehensive Survey of Honey. Editors, Chapter 5, Heinemann, London.

WHO (World Health Organization) (1997) Guidelines for predicting dietary intake of pesticide residues (revised). Global Environment monitoring System-Food Contamination and Assessment Programme (GEMS/Food) in collaboration with Codex Committee on Pesticide Residues.

